# Comparison of long read methods for sequencing and assembly of a plant genome

**DOI:** 10.1101/2020.03.16.992933

**Authors:** Valentine Murigneux, Subash Kumar Rai, Agnelo Furtado, Timothy J.C. Bruxner, Wei Tian, Qianyu Ye, Hanmin Wei, Bicheng Yang, Ivon Harliwong, Ellis Anderson, Qing Mao, Radoje Drmanac, Ou Wang, Brock A. Peters, Mengyang Xu, Pei Wu, Bruce Topp, Lachlan J.M. Coin, Robert J. Henry

## Abstract

Sequencing technologies have advanced to the point where it is possible to generate high accuracy, haplotype resolved, chromosome scale assemblies. Several long read sequencing technologies are available on the market and a growing number of algorithms have been developed over the last years to assemble the reads generated by those technologies. When starting a new genome project, it is therefore challenging to select the most cost-effective sequencing technology as well as the most appropriate software for assembly and polishing. For this reason, it is important to benchmark different approaches applied to the same sample. Here, we report a comparison of three long read sequencing technologies applied to the de novo assembly of a plant genome, *Macadamia jansenii*. We have generated sequencing data using Pacific Biosciences (Sequel I), Oxford Nanopore Technologies (PromethION) and BGI (single-tube Long Fragment Read) technologies for the same sample. Several assemblers were benchmarked in the assembly of PacBio and Nanopore reads. Results obtained from combining long read technologies or short read and long read technologies are also presented. The assemblies were compared for contiguity, accuracy and completeness as well as sequencing costs and DNA material requirements. Overall, the three long read technologies produced highly contiguous and complete genome assemblies of *Macadamia jansenii*. At the time of sequencing, the cost associated with each method was significantly different but continuous improvements in technologies have resulted in greater accuracy, increased throughput and reduced costs. We propose updating this comparison regularly with reports on significant iterations of the sequencing technologies.

## Introduction

Advances in DNA sequencing enable the rapid analysis of genomes driving biological discovery. Sequencing of complex genomes, that are very large and have a high content of repetitive sequences or many copies of similar sequences remains challenging. Many plant genomes are complex and the quality of published sequences remains relatively poor. However, improvements in long read sequencing are making it easier to generate high quality sequences for complex genomes.

We now report a comparison of three long read sequencing methods applied to the de novo sequencing of a plant, *Macadamia jansenii*. This is a rare species that is a close relative of the macadamia nut recently domesticated in Hawaii and Australia. In the wild, it grows as a multi-stemmed, evergreen tree reaching 6-9 m height with leaves having entire margins and generally in whorls of three. The nuts are small (11-16 mm diameter) and have a smooth, hard, brown shell which encloses a cream, globulose kernel that is bitter and inedible (1). The species was discovered as a single population of about 60 plants in the wild in Eastern Australia (2). This is a flowering plant (angiosperm) in the Proteaceae family that is basal to the large eudicot branch of the flowering plant phylogeny (3). The genomes of this group are poorly characterised, with most well sequenced plant genomes being either core eudicots or monocots that are plants of economic importance (4). Knowledge of the genome of this species will support efforts to conserve the endangered species in the wild and capture novel traits such a small plant stature for use in plant breeding. Sequencing of wild crop relatives is urgent as many populations are critical to diversification of crop genetics to ensure food security in response to climate change (5) but are also threatened with extinction due to changes in land use or climate (6).

Long read sequencing provides data that facilitates easier assembly of the genome than is possible with short reads (7). The length and sequence quality delivered by the available sequencing platforms has continued to improve. The reads generated can be used to assemble contigs or as a scaffold for the assembly of contigs generated with these techniques or from short reads (8). Currently, Pacific Biosciences and Oxford Nanopore Technologies are the most commonly used technologies to generate long reads. Single-molecule realtime sequencing, developed by Pacific Biosciences can generate reads in the tens of kilobases using the continuous long read sequencing mode thus enabling high-quality de novo genome assembly. Oxford Nanopore Technologies enables direct and real-time sequencing of long DNA or RNA fragments by analysing the electrical current disruption caused by the molecules as they move through a protein nanopore. More recently, BGI has introduced the single tube Long Fragment Read (stLFR) (9) technology as an alternative to the generation of real long reads. stLFR is based on DNA co-barcoding (10, 11), that is adding the same barcode sequence to sub-fragments from the original long DNA molecule. In the stLFR process, the surface of microbeads are used to create millions of miniaturized barcoding reactions in a single tube. Importantly, stLFR enables near single molecule co-barcoding by using a large excess of microbeads and a combinatorial process to make around 3.6 billion unique barcode sequences. For this reason it is expected to enable high-quality and near complete de novo assemblies. Here we compare Sequel (Pacific Biosciences), PromethION (Oxford Nanopore Technologies) and stLFR (BGI) data for the same DNA sample and evaluate the quality of the assemblies that can be generated directly from these data sets.

## Methods

### Plant material

Young leaves (40 g) of *Macadamia jansenii* were sourced from a tree with accession number 1005 and located at the Maroochy Research Facility, Department of Agriculture and Fisheries, Nambour 4560, Queensland, Australia. The specimen of *Macadamia jansenii* used in these experiments was a clonally propagated ex-situ tree planted in the arboretum at Maroochy Research Facility. None of the leaves used in these experiments were collected from wild in-situ trees. Young leaves were harvested, placed in on ice in bags and within 3 h snap frozen under liquid nitrogen and stored at −20°C until further processed for tissue pulverisation using either a mortar and pestle or the Mixer Mill as outlined below.

### Genomic DNA extraction

Leaf tissue (10 g) was first coarsely ground under liquid Nitrogen using a mortar and pestle. The mortar and pestle with the coarsely ground tissue with residual liquid nitrogen was then placed on dry ice. This step ensured the temperature of the coarsely ground tissue was maintained close to - 80°C while allowing the liquid nitrogen to evaporate off completely, an essential requirement for the pulverisation step. The coarsely ground leaf tissue was pulverised into fine powder in 50 ml steel jars using the Mixer Mill MM400 (Retsch, Germany). The pulverised leaf tissue was stored at −20°C until further required for DNA extraction. Genomic DNA (gDNA) was isolated from pulverised leaf tissue according to (12), with some modifications. Using a liquid-nitrogen cooled spatula, frozen pulverised leaf tissue (3 g) was added to 50 ml tubes (Corning or Falcon) containing warm (40°C) nuclear lysis buffer (8 ml) and 5% sarkosyl solution (5 ml). Tubes were incubated at 40°C for 45 min with periodic (every 5 min) gentle mixing by inverting the tubes. RNA was digested by adding RNase solution (10 mg/ml), the contents gently mixed by inverting the tubes followed by incubation at room temperature for 10 min. Two chloroform extractions were undertaken as follows. Chloroform (10 ml) was added to the tubes and gently mixed by inverting the tubes 50 times. The tubes were centrifuged at 3,500×g for 5 min in a swing out bucket rotor. The supernatant was transferred into fresh 50 ml tubes and the chloroform extraction repeated twice. The supernatant was transferred to fresh 50 ml tubes and the DNA precipitated using isopropanol. For every 1 ml of the supernatant, 0.6 ml of Isopropanol was added, the content gently mixed by inverting the tubes 20 to 25 times. The tubes were incubated at room temperature for 15 min and then centrifuged at 3,500×g for 5 min in a swing out bucket rotor. The supernatant was discarded and the DNA pellet was washed off any co-precipitated salts by adding 10 ml of 70% ethanol and incubating the tubes at room temperature for 30 min. The tubes were centrifuged at 3,500×g for 5 min in a swing out bucket rotor, the supernatant discarded and the DNA pellet semi dried to remove any residual 70% ethanol by incubating the tubes for 10 min upside down over filter paper. The DNA was dissolved by adding 100 μl of TE buffer and then adding incremental 50 μl of TE buffer where required. The DNA solution was transferred to 2 ml nuclease-free tubes and then centrifuged at 14,000×g for 45 min in a table top centrifuge. The supernatant was carefully transferred to fresh 2 ml tubes and the quality checked on a spectrophotometer and resolving the DNA on a 0.7% agarose gel. The DNA was then stored at −20°C until used for sequencing.

### PacBio gDNA library preparation and sequencing

DNA sequencing libraries were prepared using the Template Prep Kit 1.0-SPv3 (PacBio, 100-991-900) according to the protocol for >30 kb SMRTbell Libraries (PacBio, Part # PN 101-024-600 Version 05). Genomic DNA (15 μg) was not fragmented, and was instead just purified with AMPure PB beads. The purified gDNA (10 μg) was treated with Exonuclease VII, followed by a DNA damage repair reaction, an end-repair reaction, and purification with AMPure PB beads. Adapters were ligated to the purified, blunt-ended DNA fragments in an overnight incubation. The adapter ligated sample was digested with Exonuclease III and Exonuclease VII to remove failed ligation products, followed by purification with AMPure PB beads. The purified sample was size selected using the Blue Pippin with a dye-free, 0.75% agarose cassette and U1 marker (Sage Science, BUF7510) and the 0.75% DF Marker U1 high-pass 30-40 kb vs3 run protocol, with a BP-start cut-off of 35000 bases. After size selection, the samples were purified with AMPure PB beads, followed by another DNA damage repair reaction, and a final purification with AMPure PB beads. The final purified, size-selected library was quantified on the Qubit fluorometer using the Qubit ds-DNA HS assay kit (Invitrogen, Q32854) to assess the concentration, and a 0.4% Megabase agarose gel (BioRad, 1613108) to assess the fragment size. Sequencing was performed using the PacBio Sequel I (software/chemistry v6.0.0). The library was prepared for sequencing according to the SMRT Link sample setup calculator, following the standard protocol for Diffusion loading with AMPure PB bead purification, using Sequencing Primer v3, Sequel Binding Kit v3.0 and the Sequel DNA Internal Control v3. The polymerase-bound library was sequenced on 8 SMRT Cells with a 10 h movie time using the Sequel Sequencing Kit 3.0 (PacBio, 101-597-900) and a Sequel SMRT Cell 1M v3 (PacBio, 101-531-000). Library preparation and sequencing was performed at the Institute for Molecular Bioscience Sequencing Facility (University of Queensland).

### ONT library preparation and sequencing

The quality of the DNA sample was assessed in NanoDrop, Qubit, and the Agilent 4200 TapeStation system. The DNA sample was sequenced on the Oxford Nanopore Technologies (ONT)-MinION and PromethION. The MinION library was prepared from 1,500 ng input DNA using the ligation sequencing kit (SQK-LSK109, ONT) according to the manufacturer’s protocol except the End-repair and end-prep reaction and ligation period were increased to 30 min. Third party reagents NEBNext end repair/dA-tailing Module (E7546), NEBNext FFPE DNA Repair Mix(M6630), and NEB Quick Ligation Module (E6056) were used during library preparation. The adapters-ligated DNA sample was quantified using Qubit^®^ dsDNA HS Assay Kit (Thermofisher). The MinION flowcell R9.4.1 (FLO-MIN106, ONT) was primed according to the manufacturer’s guidelines before loading a library mix (75 μl) containing 438 ng of adapters-ligated DNA, 25.5 μl LB (SQK-LSK109, ONT), and 37.5 μl SQB (SQK-LSK109, ONT). The MinION sequencing was performed using MinKNOW (v1.15.4), and a standard 48 h run script. Before preparing the PromethION library, short DNA fragments (<10 kb) were first depleted from DNA sample (9 μg) as described in the manufacturer’s instructions for the Short Read Eliminator (SRE) kit (SKU SS-100-101-01, Circulomics Inc). The PromethION library was prepared from 1200 ng SRE-treated DNA using ligation sequencing kit (SQK-LSK109, ONT). All steps in the library preparation were the same as the MinION library preparation except the adapters-ligated DNA was eluted in 25 μl of Elution Buffer. The PromethION flowcell (FLO-PRO002) was primed according to the manufacturer’s guidelines before loading a library mix (150 μl) containing 390 ng of adapters-ligated DNA (24 μl), 75 μl of SQB and 51 μl of LB (SQK-LSK109, ONT). Sequencing was performed using MinKNOW (v3.1.23), and a standard 64 h run script. The sequencing run was stopped at 21 h and nuclease flush was performed to recover clogged pores. The Nuclease flushing mix was prepared by mixing 380 μl of Nuclease flush buffer (300 mM KCl, 2 mM CaCl2, 10 mM MgCl2, 15 mM HEPES pH 8) and 20 μl of DNase I (M0303S, NEB). The Nuclease Flushing mix was loaded into the flow cell and incubated for 30 min. The flow cell was then primed as mentioned above and loaded with the fresh library mix (150 μl) containing 390 ng of adapters-ligated DNA and rerun the standard 64 h run script using MinKNOW. Refuelling of the sequencing run was performed at each 24 h by adding 150 μl of diluted SQB (1:1, SQB:nuclease free water) to keep the stable translocation speed of sequencing. ONT fast5 reads were basecalled using Guppy v3.0.3 with the config file dna_r9.4.1_450bps_hac_prom.cfg and parameters --qscore_filtering -q 0 --recursive --device “cuda:0 cuda:1 cuda:2 cuda:3”.

### BGI library preparation and sequencing

stLFR sequencing libraries were prepared using the MGIEasy stLFR Library Prep Kit (MGI, Shenzhen, China) following the manufacturer’s protocol. Briefly, genomic DNA samples were serially diluted and then quantified using the Qubit™ dsDNA BR Assay Kit (Invitrogen, Carlsbad, CA) and the Qubit™ dsDNA HS Assay Kit (Invitrogen, Carlsbad, CA) for a more accurate quantification result. Around 1.5 ng of original genomic DNA molecules were used for library preparation. In the first step, transposons composed of a capture sequence and a transposase recognition sequence were inserted at a regular interval along the genomic DNA molecules. Next, these transposon inserted DNA molecules were hybridized with barcode labelled 3 μm diameter magnetic beads containing oligonucleotide sequences with a PCR primer annealing site, an stLFR barcode, and a sequence complementary to the capture sequence on the transposon. After hybridization, the barcode was transferred to the transposon inserted DNA sub-fragments through a ligation step. The excess oligonucleotides and transposons were then digested with exonuclease and the transposase enzyme was denatured with sodium dodecyl sulfate. Next, the second adapter was introduced by a previously described 3’-branch ligation using T4 ligase (13). Finally, PCR amplification was performed using primers annealing to the 5’ bead and 3’-branch adapter sequences. The PCR reaction was purified using Agencourt^®^ AMPure XP beads (Beckman Coulter, Brea, CA) and quantified using the Qubit™ dsDNA HS Assay Kit (Invitrogen, Carlsbad, CA). The PCR product fragment sizes were assessed using an Agilent High Sensitivity DNA Kit (Agilent, 5067-4626) on a Agilent 2100 Bioanalyzer. The average fragment size of the prepared stLFR library was 1003 bp. 20 ng of PCR product from the stLFR library was used to prepare DNA Nano Balls (DNBs) using the MGISEQ-2000RS High Throughput stLFR Sequencing Set (MGI, Shenzhen, China) following the manufacturer’s protocol. The prepared DNB library was loaded onto two lanes of a MGISEQ-2000RS flow cell (MGI, Shenzhen, China) and then sequenced on a MGISEQ-2000RS (MGI, Shenzhen, China) using the MGISEQ-2000RS stLFR sequencing Set (MGI, Shenzhen, China). Library preparation and sequencing were performed at the BGI Australia Sequencing Facility (CBCRC Level 6, Herston, QLD) and BGI-Shenzhen (Shenzhen, China).

### Illumina sequencing

Illumina library was prepared using the Nextera Flex DNA kit. The library was sequenced on an SP flow cell (14%) of the Illumina Nova Seq 6000 sequencing platform (The Ramaciotti Centre, University of New South Wales, Australia) using the paired-end protocol to produce 112 million 150 bp reads in pairs, an estimated 43× genome coverage. The median insert size was 713 bp.

### Sequence read preparation

ONT read length and quality was calculated with NanoPlot v1.22 (14). Long reads from PacBio and ONT were prepared using two or three alternative strategies respectively:

- All: no filtering of reads
- Filtered: ONT long reads were adapter-trimmed using Porechop v0.2.4 (Porechop, RRID:SCR_016967) (15). ONT and PacBio reads were filtered using Filtlong v0.2.0 (16) by removing 10% of the worst reads and reads shorter than 1 kb.
- Pass (ONT only): only the passed reads were used (average base call quality score above 7).

The PacBio subreads were randomly subsampled down to a 32× genome coverage using Rasusa v0.1.0 (17). Raw Illumina and BGI short reads were adapter-trimmed using Trimmomatic v0.36 (Trimmomatic, RRID:SCR_011848) (18) (LEADING:3 TRAILING:3 SLIDINGWINDOW:4:15 ILLUMINACLIP:2:30:10 MINLEN:36). PolyG tail trimming was performed on the Illumina reads using fastp v0.20.0 (fastp, RRID:SCR_016962) (19).

### Genome size estimation

K-mer counting using the trimmed Illumina and BGI reads was performed using Jellyfish v2.210 (Jellyfish, RRID:SCR_005491) (20) generating k-mer frequency distributions of 21-, 23- and 25-mers. The histogram of the k-mer occurrences were processed by GenomeScope (GenomeScope, RRID:SCR_017014) (21), which estimated a genome haploid size of 653 and 616 Mb with around 71% and 74% of unique content and a heterozygosity level of 0.65% and 0.77% from Illumina and BGI reads respectively.

### Assembly of genomes

De novo assembly of ONT and PacBio reads were performed using Redbean v2.5 (WTDBG, RRID:SCR_017225) (22), Flye v2.5 (Flye, RRID:SCR_017016) (23), Canu v1.8 (ONT) or v1.9 (PacBio) (Canu, RRID:SCR_015880) (24), Raven v0.0.0 (25) with default parameters. For Redbean, Flye and Canu, the estimated genome size was set to 780 Mb. For ONT data, four rounds of consensus correction were performed using Racon v1.4.6 (Racon, RRID:SCR_017642) (26) with recommended parameters (-m 8 -x -6 -g -8 - w 500) based on minimap2 v2.17-r943-dirty (27) overlaps, followed by one round of Medaka v0.8.1 (28) using the r941_prom_high model. The resulting consensus sequence was polished with Pilon v1.23 (Pilon, RRID:SCR_014731) (29) using the Illumina reads mapped with BWA-MEM v0.7.13 (BWA, RRID:SCR_010910) (30) and with the settings to fix bases. Polishing of the Medaka consensus sequence with Illumina reads was also performed by NextPolish v1.1.0 (31) with default settings (BWA for the mapping step). Hybrid assembly was generated with MaSuRCA v3.3.3 (MaSuRCA, RRID:SCR_010691) (32) using the Illumina and the ONT or PacBio reads and using Flye v2.5 to perform the final assembly of corrected mega-reads. Diploid de novo genome assembly of PacBio reads was performed with FALCON v1.3.0 (FALCON, RRID:SCR_016089) (33) using a genome size of 780 Mb, a length cutoff of 40,740 bp and a seed read coverage cutoff of 30. A total of 19 Gb of preassembled reads was generated (24× coverage). After assembly and haplotype separation by FALCON-Unzip v1.2.0 (33), polishing was performed as part of the FALCONUnzip workflow. PacBio reads were mapped to the primary FALCON-Unzip assembly using minimap2 v2.17-r954-dirty (27). A read coverage histogram was generated from this alignment using Purge Haplotigs v.1.1.0 (34) to obtain the read depth cutoff values (-l 17 -m 52 -h 190) required to identify redundant contigs. Illumina reads were assembled using SPAdes v3.13.1 (SPAdes, RRID:SCR_000131) (35).

Two lanes of stLFR reads for the same sample were demultiplexed using a sub-function of SuperPlus v1.0 (36) and combined for the downstream analysis. Adapter sequences were removed from read data using Cutadapt v2.4 (cutadapt, RRID:SCR_011841) (37) with the recommended parameters (--no-indels -O 10 --discard-trimmed -j 42). Read sequences were then converted to 10X Genomics’ format by BGI’s inhouse software, which contains three steps: 1) Change the format of reads’ head from MGI to Illumina. 2) Change the quality number of “N” base from 33 (ASIC II code = !) to 35 (ASIC II code = #) to meet the 10X Genomics’ quality system. 3) Merge two or more barcodes into one barcode randomly due to the limitation of barcode types for 10x Genomics. To meet the memory requirement of the assembler, the barcodes with less than 10 reads were removed from the dataset. De novo assembly was performed by Supernova v2.1.1 (Supernova assembler, RRID:SCR_016756) (38) using the suggested parameters (--maxreads=2100000000 --accept-extreme-coverage --nopreflight). TGS-GapCloser v1.0.0 (TGS-GapCloser, RRID:SCR_017633) (39, 40) was used to fill the gaps between contigs within same scaffolds, and this process was performed under the use of error-corrected ONT data by Canu.

### Assembly comparison

Assembly statistics were computed using QUAST v5.0.2 (QUAST, RRID:SCR_001228) (41) with a minimum contig length of 10 kb and the parameters --fragmented --large. We compared the assemblies with the published reference genome of *Macadamia integrifolia* v2 (Genbank accession: GCA_900631585.1). To estimate the base accuracy, QUAST was used to compute the number of mismatches and indels as compared to the Illumina assembly. To evaluate the completeness of the genome, the assemblies were subjected to the Benchmarking Universal Single-Copy Orthologs v3.0.2 (BUSCO, RRID:SCR_015008 (42) with the eudicotyledons_odb10 database (2121 genes).

## Results

### ONT genome assembly

For the ONT sequencing, we combined the results of one PromethION and one MinION flow cell, generating a total of 24.9 Gb of data with a read length N50 of 27.8 kb (Table 1). The PromethION flow cell and the MinION flow cell generated 23.2 Gb and 1.7 Gb of data respectively, with a read length N50 of 28.5 kb and 16.6 kb and a median read quality of 6.3 and 8.9. ONT reads were assembled using four different long-read assemblers (Redbean, Flye, Canu, Raven) and three different read subsets representing different genome coverage (21×, 28× and 32×). The statistics for each assembly are shown in Table S1. Canu and Flye generated the largest and most contiguous assemblies while Raven produced the smallest and less contiguous assembly (~720 Mb, contig N50 ~500 kb) followed by Redbean (~750 Mb, contig N50 ~700 kb). It is worth noting that Flye consistently produced assemblies of around 811 Mb with a contig N50 of approximately 1.5 Mb whereas Canu, Redbean and Raven assembly contiguity improved as the read coverage increased. In particular, the Canu contig N50 significantly improved from 706 kb (21×) to 1.43 Mb (32×). Flye was approximately five times faster than Canu.

**Table 1.**
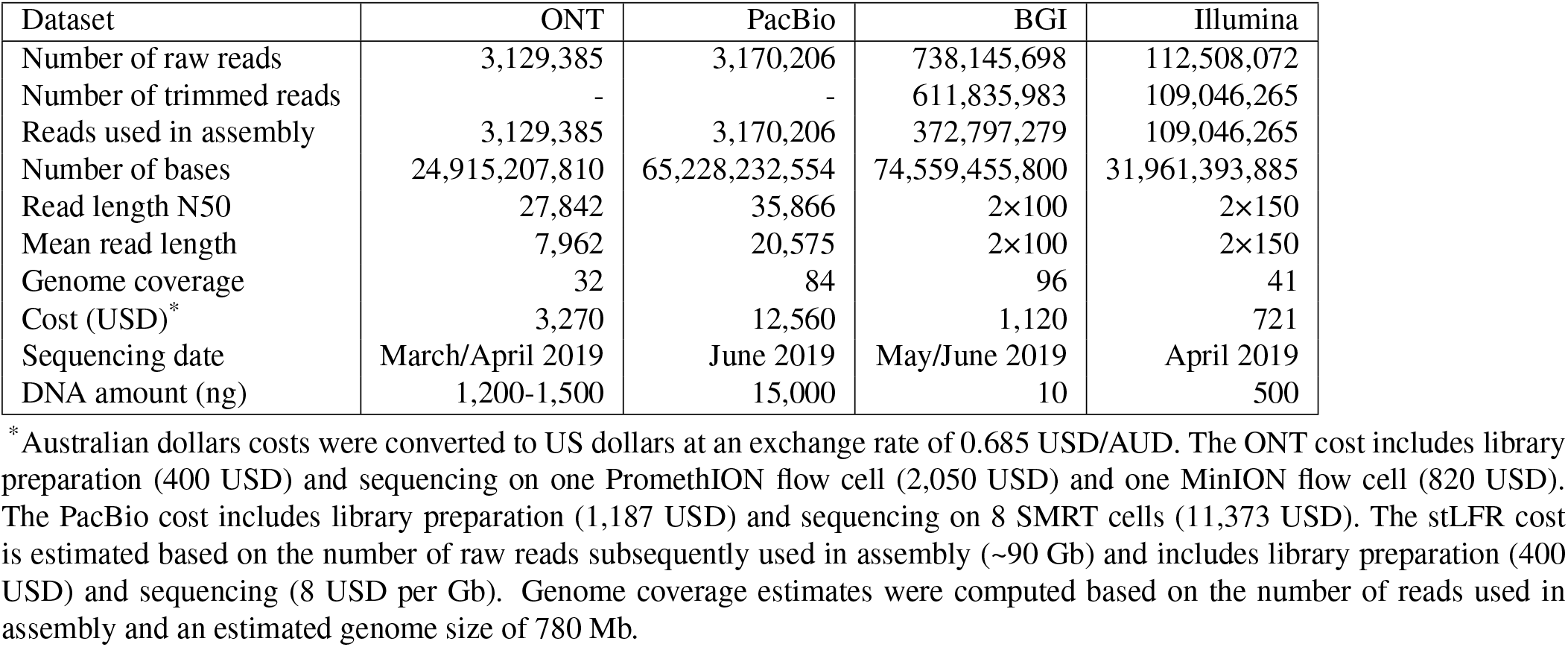
Sequencing data

We subsequently polished the Flye and Canu draft assemblies using the ONT long reads followed by the Illumina short reads. Two softwares to fix base errors using short reads were compared: the widely used tool Pilon and the recently developed algorithm NextPolish. Those polishing steps greatly improved the genome completeness as indicated by the percentage of complete BUSCO which increased from 79% (Flye) and 69% (Canu) to 88% after long-read polishing and 95% after long-read and short-read polishing (Table S2). As an estimation of the base accuracy, we computed the number of mismatches and indels as compared to the Illumina assembly. The Canu assembly was less accurate (Fig. 1) and contained a slightly higher number of duplicated genes (16-17%) as compared to the Flye assembly (13-14%) (Table S2). The base accuracy metrics showed that NextPolish performed better than Pilon. In particular, the number of indels per 100 kbp was greatly reduced after polishing with NextPolish as compared to Pilon (Flye: 43 vs 83 indels per 100 kbp, Canu: 68 vs 107 indels per 100 kbp). The genome completeness was slightly better after two iterations of NextPolish (95.5%) than after two iterations of Pilon (95.2%) (Table S1). As an alternative method to long-read-only assembly followed by polishing with short reads, we generated an hybrid assembly using MaSuRCA. The ONT + Illumina assembly showed a lower contiguity (contig N50 = 1.18 Mb) and a slightly lower accuracy but a similar completeness (94.8% complete BUSCO including 15.5% duplicated BUSCO) as the best Flye and Canu assemblies with subsequent polishing with Illumina reads (Fig. 1, Table S1 and S2).

**Fig. 1.**
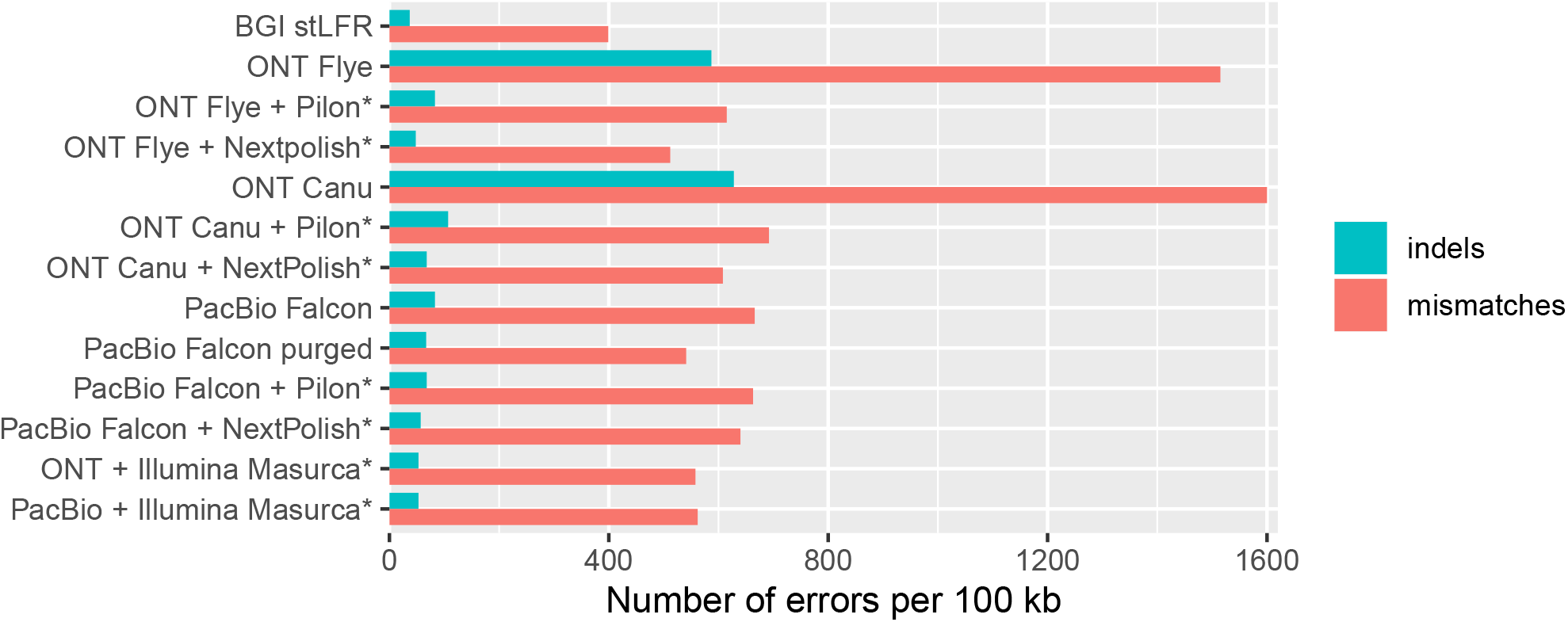
Base accuracy of assemblies as compared to Illumina assembly (*Assemblies generated or polished using Illumina reads)

### PacBio genome assembly

With eight single-molecular real-time cells in the PacBio Sequel platform, we generated 3,170,206 subreads with a read length N50 of 35.9 kb and representing a total of 65.2 Gb (Table 1). The data correspond to ~84× coverage of the estimated 780 Mb genome size. PacBio assembly was performed using the FALCON assembler, followed by phasing and polishing using FALCON-Unzip. The resulting primary assembly consisted of 1,333 contigs totaling 871 Mb in length, with half of the assembly in contigs of 1.38 Mb or longer (Table 2). FALCON-Unzip also generated 2,488 alternate haplotigs spanning 495 Mb (i.e. 57% of the genome was haplotype-resolved), with a contig N50 of 333 kb. BUSCO analysis on primary contigs showed around 26% of duplicated genes suggesting the presence of homologous primary contigs 2. The Purge Haplotigs pipeline identified 569 primary contigs representing 112 Mb as likely alternate haplotypes (Table S3). These contigs were transferred to the haplotigs set. The curated primary haploid assembly consisted of 762 contigs totaling 758 Mb with contig N50 of 1.59 Mb and contained less duplicated genes (16%) with minimal impact on genome completeness (95% complete BUSCO) (Fig. 2, Table S3). We conducted the assembly on the same PacBio data using three other long reads assemblers: Redbean, Flye and Canu (Table S4) and the hybrid assembler MaSuRCA. The Redbean assembly was the most fragmented (contig N50 = 649 kb) and the least complete (89% complete BUSCO). The Flye assembly showed a similar contiguity as the Falcon assembly (contig N50 = 1.47 Mb) but was smaller in size (767 Mb) likely due to collapsed haplotypes. The Canu assembly was much larger (1.2 Gb) but contained a higher fraction of duplication as reported by QUAST (1.64) and confirmed by the percentage of duplicated BUSCO (53%). The PacBio + Illumina hybrid assembly generated by MaSuRCA showed a slightly lower contiguity (contig N50 = 1.22 Mb), a higher accuracy and a similar completeness(94.9% complete BUSCO including 15.7% duplicated BUSCO) as the Falcon assembly with subsequent short-read polishing (Fig. 1, Table S4 and S5).

**Table 2.** Assembly statistics before short read polishing

| Dataset | ONT | PacBio | BGI |
| --- | --- | --- | --- |
| Assembler | Flye | Falcon | Supernova2 |
| Polishing | Racon, Medaka | Falcon-Unzip | - |
| Assembly size (Mb) | 808 | 871 | 752 |
| Number of contigs | 2,243 | 1,333 | 19,954 |
| Number of scaffolds | - | - | 5,065 |
| Contig N50 (Mb) | 1.49 | 1.38 | 0.036 |
| Scaffold N50 (Mb) | - | - | 3.54 |
| Contig NG50 (Mb) | 1.77 | 1.61 | 0.028 |
| Scaffold NG50 (Mb) | - | - | 3.63 |
| Contig L50 | 105 | 173 | 4,942 |
| Longest contig (Mb) | 14.23 | 10.54 | 0.52 |
| Longest scaffold (Mb) | - | - | 30.14 |
| Genome fraction (%) | 32.9 | 56.1 | 50.5 |
| Complete BUSCO (%) | 88.4 | 95.5 | 88.3 |

QUAST analysis was performed using a minimum contig size of 10 kb.

**Fig. 2.**
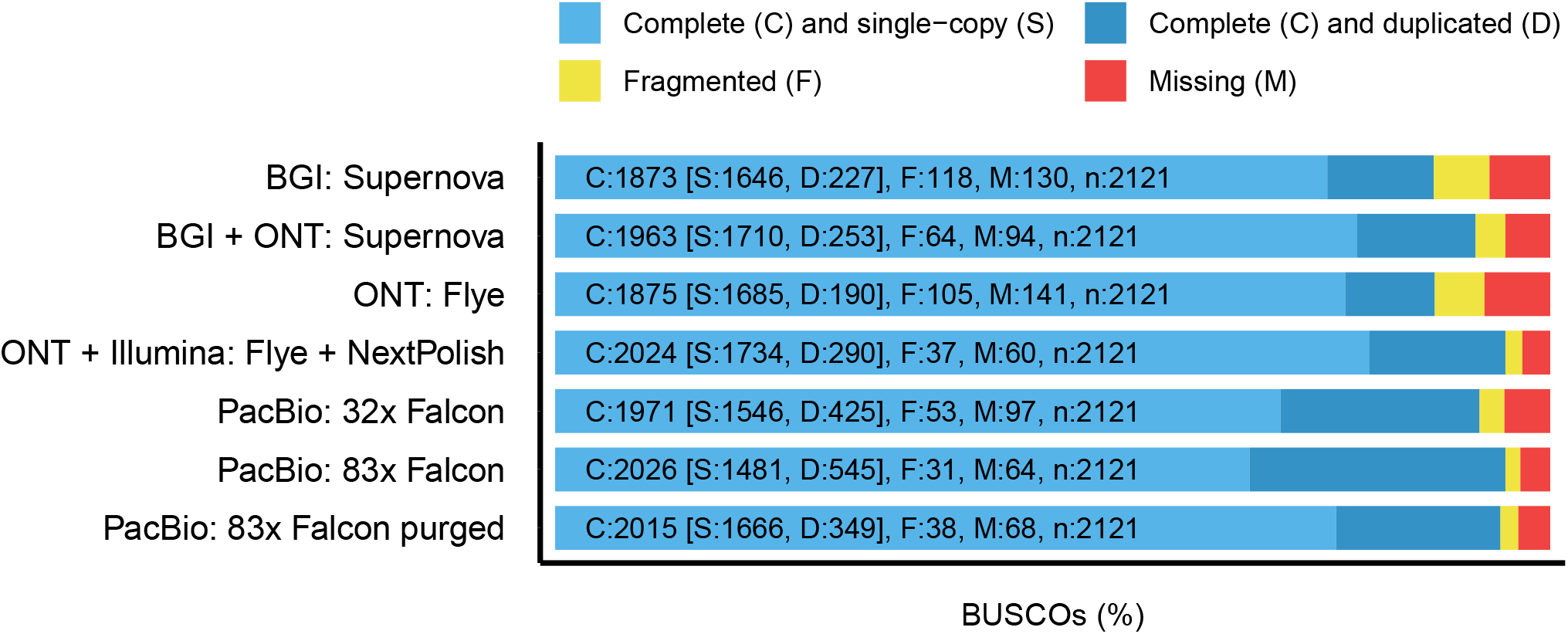
BUSCO analysis of assemblies using the eudicotyledons dataset. The x-axis depicts the percentage of complete and single-copy, complete and duplicated, fragmented and missing BUSCO and the y-axis indicates the assembly assessed.

Using a quality filtered subset of the subreads (equivalent to ~67× genome coverage) led to a slight improvement in the assembly contiguity (Redbean) without impacting on the genome completeness (only Rebean and Flye were tested due to high computational requirements of Canu and Falcon) (Table S4). We next investigated the effects of genome coverage on the assembly quality by using approximately half of the subreads. Two subsets of reads corresponding to 4 SMRT cells and equivalent to a 43× and 39× coverage were assembled using Flye. The assembly size slightly decreased from 767 Mb to 765 Mb and 762 Mb respectively and became a bit more fragmented with the contig N50 decreasing from 1.47 Mb to 1.41 Mb and 1.28 Mb but with no impact on the genome completeness (94.3% and 94.7% of complete BUSCO as compared to 94.5%) (Table S4). Finally, in order to compare PacBio and ONT technologies, we randomly subsampled the PacBio subreads down to a coverage equivalent to the ONT data (32×). The resulting Flye assembly showed a similar size of 764 Mb, a lower contiguity (contig N50 = 1.26 Mb) and a similar genome completeness (94.7% complete BUSCO) as the 83× coverage assembly (Table S4). The Falcon assembly was the most affected by the coverage drop with a decrease in the contig N50 from 1.38 Mb to 684 kb. Canu was also quite robust to the coverage drop with a decrease in contig N50 from 2.07 Mb to 1.65 Mb.

### stLFR genome assembly

stLFR generated 738 million 100 bp paired-end reads. To meet the requirements of the assembler, the barcodes with less than 10 reads were removed which resulted in 373 million reads representing 74.6 Gb of data and corresponding to approximately 96× coverage of the genome (Table 1). stLFR reads were assembled using Supernova2 into an assembly of 40,789 scaffolds totaling 880 Mb in length. 5,065 scaffolds were larger than 10 kb with a total length of 752 Mb and a N50 of 3.54 Mb for scaffold and 35.6 kb for contig (Table 2). The stLFR assembly was the most accurate with the lowest number of mismatches and indels identified as compared to the Illumina assembly (Fig. 1). Conserved BUSCO gene analysis revealed that the stLFR assembly contained 88.3% of complete genes from the eudicotyledons dataset (Fig. 2). Inclusion of ONT data to fill the gaps within scaffolds led to a 29-fold increase in the contig N50 length from 35.6 kb to 1.05 Mb and a 11-fold decrease in the number of gaps from 24,933 to 2,284 (Table 3). The scaffold N50 slightly dropped by 0.02 Mb due to the adjustment of the estimated gaps. The total length also increased correspondingly to 894 Mb and to 767 Mb for scaffolds larger than 10 kb. The largest contig size increased from 518 kb to 9.7 Mb. In addition, the genome completeness was improved in the gap-filled assembly, with BUSCO detecting 4.8% more complete genes (Fig. 2).

**Table 3.**
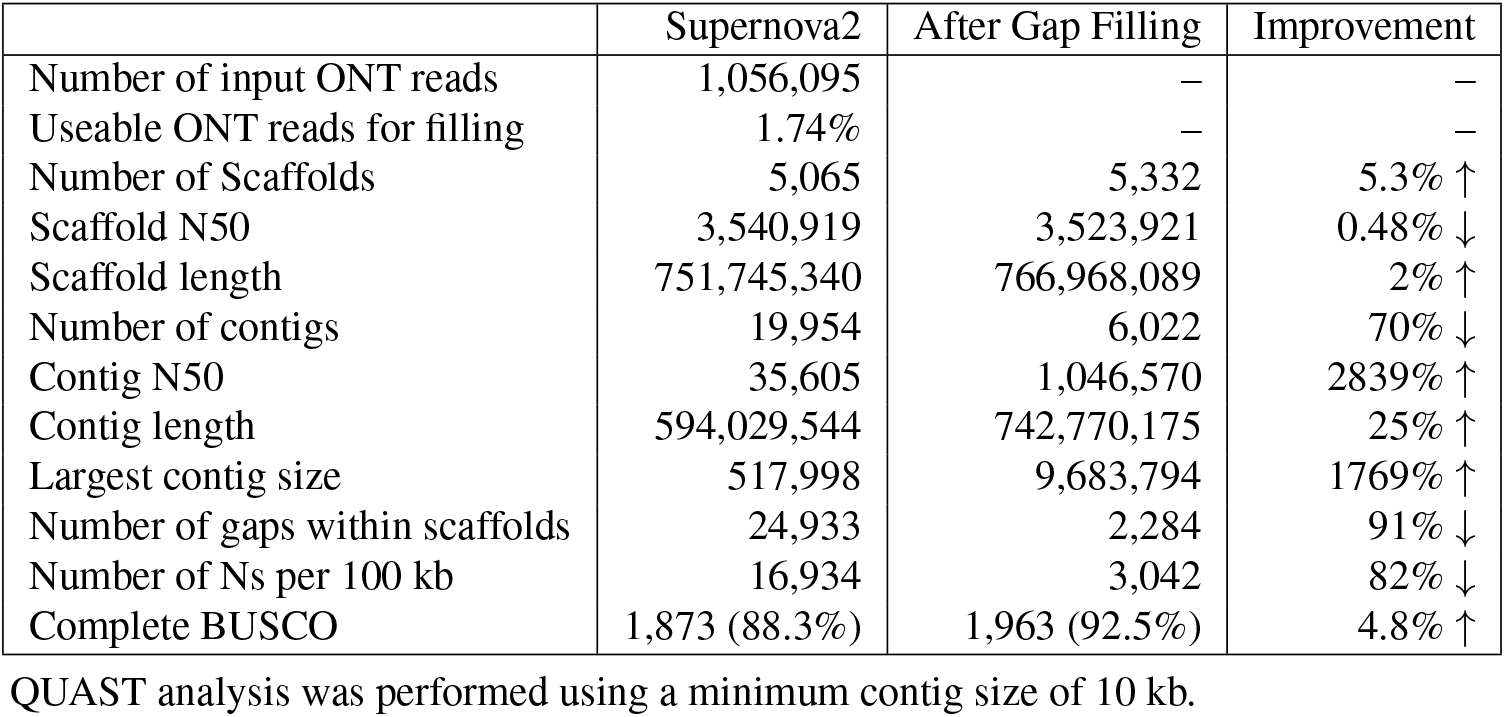
Gap filling for stLFR assembly using error-corrected ONT reads

## Discussion

We report a comparison of three long-read sequencing datasets generated from the same plant DNA sample. *M. jansenii* was selected for this study because of its significance in conservation and breeding. All four species of *Macadamia* are listed as threatened under Australian legislation but *M. jansenii* is particularly vulnerable given it has been recorded at only one location. *M. jansenii* has not been domesticated and its small and bitter nuts are obstacles that restrict simple introgression in breeding. However, the characteristic small tree size, being 50% smaller than commercial cultivars, is of interest for use in high-density orchard design and it is being trialled as a rootstock for this purpose (43). It is the most northern *Macadamia* species and may be a source of genes for adaptation to warmer climates (44). Hybrids of *M. integrifolia* and *M. jansenii* have been produced.

The amount of sequencing data produced by each platform corresponds to approximately 84× (PacBio Sequel), 32× (ONT) and 96× (BGI stLFR) coverage of the macadamia genome. The cost of generating 1 Gb of sequencing data (including the library preparation) was 193 USD for PacBio Sequel I, 97 USD for ONT PromethION and 12 USD for BGI stLFR (raw reads subsequently used in assembly). Virtual long reads were generated using the stLFR protocol. This technology benefits from the accuracy and the low cost of a short-read sequencing platform while providing long range information. It was the cheapest and most accurate as it generated an assembly with the fewest single base and indel errors. Furthermore, the assembly generated by Supernova was phased. That said, the stLFR assembly was more fragmented than the others. We also demonstrated that stLFR could be used as a complementary technology to ONT. Indeed, the inclusion of Nanopore reads significantly increased the stlFR assembly contiguity with a N50 reaching 1 Mb and improved the genome completeness. When all the reads were incorporated, the assemblies generated using the PacBio and ONT data were comparable in terms of assembly contiguity (contig N50 of around 1.5 Mb) and genome completeness (95% of complete BUSCO). However, when we utilised the same amount of data for each platform (32× coverage), the contiguity of the PacBio assembly produced by Falcon was halved and became only half the size of the ones from the ONT Flye or Canu assemblies. The Flye assembler proved to be more robust to the PacBio coverage drop as the assembly contig N50 only dropped to 1.26 Mb. Additionally, we found that polishing the ONT assembly with Illumina short reads was required to reach a similar genome completeness to that of the PacBio assembly. For both ONT and PacBio data, the highest assembly contiguity was obtained with a long-read only assembler as compared to an hybrid assembler incorporating both the short and long reads.

Since the sequence data was generated, the PacBio SMRT platform has transitioned from the Sequel I to the Sequel II instrument, with a 8-fold increase in the data yield. The latest platform produces high-fidelity reads that are more accurate than the continuous long reads assembled in this study. Consequently the cost to generate a similar PacBio assembly on the Sequel II system will be dramatically reduced and the assembly quality is likely to be improved while requiring less computational resources.

The DNA material requirements to prepare the sequencing library is another important parameter to consider when choosing a sequencing technology. For ONT sequencing, it is recommended to obtain at least 1-2 μg of high molecular weight DNA. The stLFR library construction requires at least 10 ng of high molecular weight DNA. PacBio SMRT sequencing has a high genomic DNA input requirements of 5-20 μg of high molecular weight DNA for standard library protocol depending on the genome size but the PacBio low DNA input protocol has reduced this requirement to as low as 100 ng per 1 Gb genome size (45). Furthermore, PacBio recently released an amplification-based ultra-low DNA input protocol starting with 5 ng of high molecular weight DNA.

The computational requirements should be considered and will largely depend on the genome size of the species of interest. There were important differences in the assembly run time and memory usage depending on the tool used. For instance, a large memory computing cluster of 1 TB was required to assemble the reads using Supernova or polish the assembly using Pilon. GPU accelerated computing greatly reduced the computing time for some tools such as Racon, Medaka or Raven. There are also challenges associated with the rapid evolution of technologies and softwares. For example we sometimes observed a significant improvement in the ONT assembly contiguity depending on the basecaller or assembler version used.

The three technologies produced highly contiguous and complete genome assemblies. Next, long-range scaffolding approaches such as chromosome conformation capture (Hi-C, Chicago) or physical maps technologies (optical map, restriction map) are required to order and orient the assembled contigs into chromosome-length scaffolds (46).

## Supporting information

Supplemental Datatse

## Availability of supporting data and materials

BGI, PacBio, ONT and Illumina sequencing data generated in this study have been deposited in the Sequence Read Archive under BioProject PRJNA609013 and BioSample SAMN14217788. Accession numbers are as follows: BGI (SRR11191908), PacBio (SRR11191909), ONT PromethION (SRR11191910), ONT MinION (SRR11191911) and Illumina (SRR11191912).

## Additional Files

**Table S1:** ONT genome assembly statistics using Redbean, Flye, Canu, Raven and MaSuRCA

**Table S2:** BUSCO genome completeness assessment of ONT long-read assemblies and hybrid assembly

**Table S3:** PacBio genome assembly statistics (QUAST) and genome completeness assessment (BUSCO) before and after Purge Haplotigs

**Table S4:** PacBio genome assembly statistics using Redbean, Flye, Falcon, Canu and MaSuRCA

**Table S5:** BUSCO genome completeness assessment of PacBio long-read assemblies and hybrid assembly

## List of abbreviations

AUD: Australian dollars
bp: base pairs
BGI: Beijing Genomics Institute
BUSCO: Benchmarking Universal Single-Copy Orthologs
BWA: Burrows-Wheeler Aligner
g: gram
Gb: gigabase pairs
kb: kilobase pairs
Mb: megabase pairs
mg: milligram
μl: microlitre
ml: millilitre
mm: millimeter
ng: nanogram
ONT: Oxford Nanopore Technologies
PacBio: Pacific Biosciences
QUAST: QUality ASsessment Tool
SMRT: single-molecule real-time
SPAdes: St. Petersburg genome assembler
stLFR: Single Tube Long Fragment Reads
TB: Terabyte
USD: United States Dollar

## Author’s Contributions

A.F. prepared the sample. B.T. supervised plant collection. S.K.R. performed ONT library preparation and sequencing. T.J.C.B. performed PacBio library preparation and sequencing. V.M. performed ONT and PacBio assemblies and assembly evaluation. Q.Y. and H.W. performed stLFR library preparation and sequencing. I.H supervised and reviewed stLFR library preparation and sequencing. W.T. performed stLFR assembly, gap filling and statistics for stLFR. E.A., Q.M., R.D., O.W., and B.A.P. designed stLFR experiments and performed stLFR analyses. V.M. wrote the manuscript with input from all authors. R.J.H. and L.J.M.C. designed and supervised the project.

## Acknowledgements

We acknowledge Doug Stetner and Thom Cuddihy for help with the Falcon software, Nicholas Rhodes and Chenxi Zhou for help with the MaSuRCA software, Tania Duarte for running the DNA sample in tapestation and Mobashwer Alam for provision of the Macadamia tissue samples.

## Bibliography

1. Cl Gross and Ph Weston. *Macadamia jansenii* (Proteaceae), a new species from central Queensland. Australian Systematic Botany, 5(6):725–728, 1992. ISSN 1030-1887. doi: 10.1071/SB9920725.

2. The four macadamias.http://www.wildmacadamias.org.au/the-four-macadamias. Accessed February 14, 2020.

3. M.W. Chase. Relationships between the families of flowering plants. In: Henry RJ, (ed.), Plant Diversity and Evolution: Genotypic and Phenotypic Variation in Higher Plants. CABI Pub, Wallingford, Oxfordshire, UK; Cambridge, MA, 2005. ISBN 978-0-85199-904-3.

4. Marta Brozynska, Agnelo Furtado, and Robert J. Henry. Genomics of crop wild relatives: expanding the gene pool for crop improvement. Plant Biotechnology Journal, 14(4):1070– 1085, April 2016. ISSN 14677644. doi: 10.1111/pbi.12454.

5. Michael Abberton, Jacqueline Batley, Alison Bentley, John Bryant, Hongwei Cai, James Cockram, Antonio Costa de Oliveira, Leland J. Cseke, Hannes Dempewolf, Ciro De Pace, David Edwards, Paul Gepts, Andy Greenland, Anthony E. Hall, Robert Henry, Kiyosumi Hori, Glenn Thomas Howe, Stephen Hughes, Mike Humphreys, David Lightfoot, Athole Marshall, Sean Mayes, Henry T. Nguyen, Francis C. Ogbonnaya, Rodomiro Ortiz, Andrew H. Paterson, Roberto Tuberosa, Babu Valliyodan, Rajeev K. Varshney, and Masahiro Yano. Global agricultural intensification during climate change: a role for genomics. Plant Biotechnology Journal, 14(4):1095–1098, April 2016. ISSN 14677644. doi: 10.1111/pbi.12467.

6. Robert J Henry. Innovations in plant genetics adapting agriculture to climate change. Current Opinion in Plant Biology, 13:1–6, December 2019. ISSN 13695266. doi: 10.1016/j.pbi.2019.11.004.

7. Pirita Paajanen, George Kettleborough, Elena López-Girona, Michael Giolai, Darren Heavens, David Baker, Ashleigh Lister, Fiorella Cugliandolo, Gail Wilde, Ingo Hein, Iain Macaulay, Glenn J. Bryan, and Matthew D. Clark. A critical comparison of technologies for a plant genome sequencing project. GigaScience, 8(3), 2019. ISSN 2047-217X. doi: 10.1093/gigascience/giy163.

8. Hyungtaek Jung, Christopher Winefield, Aureliano Bombarely, Peter Prentis, and Peter Waterhouse. Tools and Strategies for Long-Read Sequencing and De Novo Assembly of Plant Genomes. Trends in Plant Science, 24(8):700–724, August 2019. ISSN 13601385. doi: 10.1016/j.tplants.2019.05.003.

9. Ou Wang, Robert Chin, Xiaofang Cheng, Michelle Ka Yan Wu, Qing Mao, Jingbo Tang, Yuhui Sun, Ellis Anderson, Han K. Lam, Dan Chen, Yujun Zhou, Linying Wang, Fei Fan, Yan Zou, Yinlong Xie, Rebecca Yu Zhang, Snezana Drmanac, Darlene Nguyen, Chongjun Xu, Christian Villarosa, Scott Gablenz, Nina Barua, Staci Nguyen, Wenlan Tian, Jia Sophie Liu, Jingwan Wang, Xiao Liu, Xiaojuan Qi, Ao Chen, He Wang, Yuliang Dong, Wenwei Zhang, Andrei Alexeev, Huanming Yang, Jian Wang, Karsten Kristiansen, Xun Xu, Radoje Drmanac, and Brock A. Peters. Efficient and unique cobarcoding of second-generation sequencing reads from long DNA molecules enabling cost-effective and accurate sequencing, haplotyping, and de novo assembly. Genome Research, 29(5):798–808, May 2019. ISSN 1088-9051, 1549-5469. doi: 10.1101/gr.245126.118.

10. Radoje Drmanac. Nucleic Acid Analysis by Random Mixtures of Non-Overlapping Fragments. patent wo 2006/138284, 2006.

11. Brock A. Peters, Jia Liu, and Radoje Drmanac. Co-barcoded sequence reads from long DNA fragments: a cost-effective solution for “perfect genome” sequencing. Frontiers in Genetics, 5:466, 2014. ISSN 1664-8021. doi: 10.3389/fgene.2014.00466.

12. Agnelo Furtado. DNA extraction from vegetative tissue for next-generation sequencing. Methods in Molecular Biology (Clifton, N.J.), 1099:1–5, 2014. ISSN 1940-6029. doi: 10.1007/978-1-62703-715-0_1.

13. Lin Wang, Yang Xi, Wenwei Zhang, Weimao Wang, Hanjie Shen, Xiaojue Wang, Xia Zhao, Andrei Alexeev, Brock A. Peters, Alayna Albert, Xu Xu, Han Ren, Ou Wang, Killeen Kirkconnell, Helena Perazich, Sonya Clark, Evan Hurowitz, Ao Chen, Xun Xu, Radoje Drmanac, and Yuan Jiang. 3’ Branch ligation: a novel method to ligate non-complementary DNA to recessed or internal 3’OH ends in DNA or RNA. DNA research: an international journal for rapid publication of reports on genes and genomes, 26(1):45–53, February 2019. ISSN 1756-1663. doi: 10.1093/dnares/dsy037.

14. Wouter De Coster, Svenn D’Hert, Darrin T Schultz, Marc Cruts, and Christine Van Broeckhoven. NanoPack: visualizing and processing long-read sequencing data. Bioinformatics, 34(15):2666–2669, August 2018. ISSN 1367-4803, 1460-2059. doi: 10.1093/bioinformatics/bty149.

15. Porechop.https://github.com/rrwick/Porechop. Accessed May 23, 2019.

16. Filtlong.https://github.com/rrwick/Filtlong. Accessed May 23, 2019.

17. Rasusa.https://github.com/mbhall88/rasusa. Accessed November 11, 2019.

18. Anthony M. Bolger, Marc Lohse, and Bjoern Usadel. Trimmomatic: a flexible trimmer for Illumina sequence data. Bioinformatics (Oxford, England), 30(15):2114–2120, August 2014. ISSN 1367-4811. doi: 10.1093/bioinformatics/btu170.

19. Shifu Chen, Yanqing Zhou, Yaru Chen, and Jia Gu. fastp: an ultra-fast all-in-one FASTQ preprocessor. Bioinformatics (Oxford, England), 34(17):i884–i890, 2018. ISSN 1367-4811. doi: 10.1093/bioinformatics/bty560.

20. Guillaume Marçais and Carl Kingsford. A fast, lock-free approach for efficient parallel counting of occurrences of k-mers. Bioinformatics, 27(6):764–770, March 2011. ISSN 1460-2059, 1367-4803. doi: 10.1093/bioinformatics/btr011.

21. Gregory W Vurture, Fritz J Sedlazeck, Maria Nattestad, Charles J Underwood, Han Fang, James Gurtowski, and Michael C Schatz. GenomeScope: fast reference-free genome profiling from short reads. Bioinformatics, 33(14):2202–2204, July 2017. ISSN 1367-4803, 1460-2059. doi: 10.1093/bioinformatics/btx153.

22. Jue Ruan and Heng Li. Fast and accurate long-read assembly with wtdbg2. Nature Methods, 17(2):155–158, February 2020. ISSN 1548-7105. doi: 10.1038/s41592-019-0669-3.

23. Mikhail Kolmogorov, Jeffrey Yuan, Yu Lin, and Pavel A. Pevzner. Assembly of long, error-prone reads using repeat graphs. Nature Biotechnology, 37(5):540–546, 2019. ISSN 1546-1696. doi: 10.1038/s41587-019-0072-8.

24. Sergey Koren, Brian P. Walenz, Konstantin Berlin, Jason R. Miller, Nicholas H. Bergman, and Adam M. Phillippy. Canu: scalable and accurate long-read assembly via adaptive *k* - mer weighting and repeat separation. Genome Research, 27(5):722–736, May 2017. ISSN 1088-9051, 1549-5469. doi: 10.1101/gr.215087.116.

25. Raven.https://github.com/lbcb-sci/raven. Accessed September 13, 2019.

26. Robert Vaser, Ivan Sović, Niranjan Nagarajan, and Mile Šikić. Fast and accurate de novo genome assembly from long uncorrected reads. Genome Research, 27(5):737–746, May 2017. ISSN 1088-9051, 1549-5469. doi: 10.1101/gr.214270.116.

27. Heng Li. Minimap2: pairwise alignment for nucleotide sequences. Bioinformatics, 34(18): 3094–3100, September 2018. ISSN 1367-4803, 1460-2059. doi: 10.1093/bioinformatics/bty191.

28. Medaka. https://github.com/nanoporetech/medaka. Accessed September 5, 2019.

29. Bruce J. Walker, Thomas Abeel, Terrance Shea, Margaret Priest, Amr Abouelliel, Sharadha Sakthikumar, Christina A. Cuomo, Qiandong Zeng, Jennifer Wortman, Sarah K. Young, and Ashlee M. Earl. Pilon: An Integrated Tool for Comprehensive Microbial Variant Detection and Genome Assembly Improvement. PLoS ONE, 9(11):e112963, November 2014. ISSN 1932-6203. doi: 10.1371/journal.pone.0112963.

30. Heng Li. Aligning sequence reads, clone sequences and assembly contigs with BWA-MEM. arXiv:1303.3997[q-bio], May 2013. arXiv: 1303.3997.

31. Jiang Hu, Junpeng Fan, Zongyi Sun, and Shanlin Liu. NextPolish: a fast and efficient genome polishing tool for long read assembly. Bioinformatics (Oxford, England), November 2019. ISSN 1367-4811. doi: 10.1093/bioinformatics/btz891.

32. Aleksey V. Zimin, Guillaume Marçais, Daniela Puiu, Michael Roberts, Steven L. Salzberg, and James A. Yorke. The MaSuRCA genome assembler. Bioinformatics (Oxford, England), 29(21):2669–2677, November 2013. ISSN 1367-4811. doi: 10.1093/bioinformatics/btt476.

33. Chen-Shan Chin, Paul Peluso, Fritz J. Sedlazeck, Maria Nattestad, Gregory T. Concepcion, Alicia Clum, Christopher Dunn, Ronan O’Malley, Rosa Figueroa-Balderas, Abraham Morales-Cruz, Grant R. Cramer, Massimo Delledonne, Chongyuan Luo, Joseph R. Ecker, Dario Cantu, David R. Rank, and Michael C. Schatz. Phased diploid genome assembly with single-molecule real-time sequencing. Nature Methods, 13(12):1050–1054, December 2016. ISSN 1548-7105. doi: 10.1038/nmeth.4035.

34. Michael J. Roach, Simon A. Schmidt, and Anthony R. Borneman. Purge Haplotigs: allelic contig reassignment for third-gen diploid genome assemblies. BMC bioinformatics, 19(1): 460, November 2018. ISSN 1471-2105. doi: 10.1186/s12859-018-2485-7.

35. Anton Bankevich, Sergey Nurk, Dmitry Antipov, Alexey A. Gurevich, Mikhail Dvorkin, Alexander S. Kulikov, Valery M. Lesin, Sergey I. Nikolenko, Son Pham, Andrey D. Prjibelski, Alexey V. Pyshkin, Alexander V. Sirotkin, Nikolay Vyahhi, Glenn Tesler, Max A. Alekseyev, and Pavel A. Pevzner. SPAdes: a new genome assembly algorithm and its applications to single-cell sequencing. Journal of Computational Biology: A Journal of Computational Molecular Cell Biology, 19(5):455–477, May 2012. ISSN 1557-8666. doi: 10.1089/cmb.2012.0021.

36. Superplus split_barcode. https://github.com/MGI-tech-bioinformatics/SuperPlus/blob/master/split_barcode/split_barcode_PEXXX_42_unsort_reads.pl. Accessed August 19, 2019.

37. Marcel Martin. Cutadapt removes adapter sequences from high-throughput sequencing reads. EMBnet.journal, 17(1):10, May 2011. ISSN 2226-6089. doi: 10.14806/ej.17.1.200.

38. Neil I. Weisenfeld, Vijay Kumar, Preyas Shah, Deanna M. Church, and David B. Jaffe. Direct determination of diploid genome sequences. Genome Research, 27(5):757–767, May 2017. ISSN 1088-9051, 1549-5469. doi: 10.1101/gr.214874.116.

39. Mengyang Xu, Lidong Guo, Shengqiang Gu, Ou Wang, Rui Zhang, Guangyi Fan, Xun Xu, Li Deng, and Xin Liu. TGS-GapCloser: fast and accurately passing through the Bermuda in large genome using error-prone third-generation long reads. preprint, bioRxiv, November 2019.

40. Tgs-gapcloser.https://github.com/BGI-Qingdao/TGS-GapCloser. Accessed October 15, 2019.

41. Alexey Gurevich, Vladislav Saveliev, Nikolay Vyahhi, and Glenn Tesler. QUAST: quality assessment tool for genome assemblies. Bioinformatics, 29(8):1072–1075, April 2013. ISSN 1460-2059, 1367-4803. doi: 10.1093/bioinformatics/btt086.

42. Felipe A. Simão, Robert M. Waterhouse, Panagiotis Ioannidis, Evgenia V. Kriventseva, and Evgeny M. Zdobnov. BUSCO: assessing genome assembly and annotation completeness with single-copy orthologs. Bioinformatics, 31(19):3210–3212, October 2015. ISSN 1367-4803, 1460-2059. doi: 10.1093/bioinformatics/btv351.

43. M.M. Alam, J. Wilkie, and B.L. Topp. Early growth and graft success in macadamia seedling and cutting rootstocks. Acta Horticulturae, (1205):637–644, June 2018. ISSN 0567-7572, 2406-6168. doi: 10.17660/ActaHortic.2018.1205.79.

44. Bruce L. Topp, Catherine J. Nock, Craig M. Hardner, Mobashwer Alam, and Katie M. O’Connor. Macadamia (Macadamia spp.) Breeding. In Jameel M. Al-Khayri, Shri Mohan Jain, and Dennis V. Johnson, editors, Advances in Plant Breeding Strategies: Nut and Beverage Crops, pages 221–251. Springer International Publishing, Cham, 2019. ISBN 978-3-030-23111-8 978-3-030-23112-5. doi: 10.1007/978-3-030-23112-5_7.

45. Sarah Kingan, Haynes Heaton, Juliana Cudini, Christine Lambert, Primo Baybayan, Brendan Galvin, Richard Durbin, Jonas Korlach, and Mara Lawniczak. A High-Quality De novo Genome Assembly from a Single Mosquito Using PacBio Sequencing. Genes, 10(1):62, January 2019. ISSN 2073-4425. doi: 10.3390/genes10010062.

46. Jay Ghurye and Mihai Pop. Modern technologies and algorithms for scaffolding assembled genomes. PLoS computational biology, 15(6):e1006994, 2019. ISSN 1553-7358. doi: 10.1371/journal.pcbi.1006994.

